# Protein stability models fail to capture epistatic interactions of double point mutations

**DOI:** 10.1101/2024.08.20.608844

**Authors:** Henry Dieckhaus, Brian Kuhlman

**Affiliations:** Department of Biochemistry and Biophysics, University of North Carolina School of Medicine, Chapel Hill, North Carolina, USA; Division of Chemical Biology and Medicinal Chemistry, University of North Carolina Eshelman School of Pharmacy, Chapel Hill, North Carolina, USA; Department of Bioinformatics and Computational Biology, University of North Carolina School of Medicine, Chapel Hill, North Carolina, USA; Lineberger Comprehensive Cancer Center, University of North Carolina School of Medicine, Chapel Hill, North Carolina, USA

**Keywords:** Protein stability, epistasis, point mutations, deep learning, protein design

## Abstract

There is strong interest in accurate methods for predicting changes in protein stability resulting from amino acid mutations to the protein sequence. Recombinant proteins must often be stabilized to be used as therapeutics or reagents, and destabilizing mutations are implicated in a variety of diseases. Due to increased data availability and improved modeling techniques, recent studies have shown advancements in predicting changes in protein stability when a single point mutation is made. Less focus has been directed toward predicting changes in protein stability when there are two or more mutations, despite the significance of mutation clusters for disease pathways and protein design studies. Here, we analyze the largest available dataset of double point mutation stability and benchmark several widely used protein stability models on this and other datasets. We identify a blind spot in how predictors are typically evaluated on multiple mutations, finding that, contrary to assumptions in the field, current stability models are unable to consistently capture epistatic interactions between double mutations. We observe one notable deviation from this trend, which is that epistasis-aware models provide marginally better predictions on stabilizing double point mutations. We develop an extension of the ThermoMPNN framework for double mutant modeling as well as a novel data augmentation scheme which mitigates some of the limitations in available datasets. Collectively, our findings indicate that current protein stability models fail to capture the nuanced epistatic interactions between concurrent mutations due to several factors, including training dataset limitations and insufficient model sensitivity.

**Significance:** Protein stability is governed in part by epistatic interactions between energetically coupled residues. Prediction of these couplings represents the next frontier in protein stability modeling. In this work, we benchmark protein stability models on a large dataset of double point mutations and identify previously overlooked limitations in model design and evaluation. We also introduce several new strategies to improve modeling of epistatic couplings between protein point mutations.

## Introduction

Thermodynamic stability is an important property that can impact the fitness of a protein^1,2^. Molecular biologists often introduce mutations to probe structure-function relationships within proteins, and aberrant stability profiles are implicated in a variety of diseases^3,4^. Additionally, as engineered proteins are increasingly used as therapeutics^5^ and research tools^6^, their stability must often be optimized to improve production yields and efficacy^7^. Recent advancements in assay design and transfer learning have enabled deep neural networks to predict the change in stability (ΔG) caused by single point mutations faster and more accurately than prior approaches^8–10^. However, relatively few studies have attempted to explicitly model multiple point mutations, for a few reasons. Not only is reliable stability data less abundant for multiple mutations, but the possible mutation space also increases exponentially with the number of mutations, resulting in a sparse energy landscape that is difficult to model.

In this study, we focus on the task of predicting changes in stability (ΔΔG) caused by double point mutations. We are partially motivated by the observation that protein engineers often seek to identify clusters of two or more mutations which may improve stability beyond levels achievable with single mutant sweeps through favorable couplings such as hydrogen bonding or apolar packing^11^. Also, biological researchers must sometimes contend with multiple concurrent mutations introduced by cancer^12^, bacteria^13^, or viruses^14^. A single mutant stability model can be used to approximate double mutant ΔΔG by simply adding the two constituent single mutant contributions. The drawback of this additive approach is that it omits any epistatic coupling that may arise from the interaction of the two mutations. As such, the utility of a double mutant predictor is derived from its ability to provide improvements relative to the additive predictions provided from its equivalent single mutant model. Despite this observation, double mutant stability models are rarely evaluated in this way. We posit that this represents a significant blind spot in our current understanding of protein stability models, which we aim to address in this work.

To that end, we develop a novel method for modeling stability changes due to double point mutations which we call ThermoMPNN-Double (“ThermoMPNN-D”). We analyze the largest available double mutant dataset and introduce a new data augmentation protocol to address shortcomings in data availability. We evaluate ThermoMPNN-D against popular methods from the literature, and we take the additional step of evaluating each predictor against its own additive equivalent. We show that ThermoMPNN-D and its single mutant analogue, ThermoMPNN, provide competitive performance on two datasets of double mutants gathered on a diverse set of proteins. We use deep mutational scanning (DMS) data as an orthogonal test set, finding that the methods Mutate Everything and FoldX perform the best on this task. Overall, we find that epistasis-aware double mutant models rarely outperform their single mutant counterparts, with the notable exception that they provide improved prediction of stabilizing double mutants.

## Results

### Adapting ThermoMPNN to model double mutations

We developed a novel neural network, ThermoMPNN-D, to model double point mutations by making several modifications to the previously described ThermoMPNN framework^10^ (Fig. 1A). ThermoMPNN is a structure-based protein stability model that extracts learned residue embeddings from ProteinMPNN and passes these features through a lightweight prediction head to obtain single mutant ΔΔG predictions. ProteinMPNN is a graph neural network trained to predict protein sequences from the 3D structure of the protein^15^. Both models use message passing to encode the local context surrounding the residue of interest, including the relative positions of nearby residues. In this way, they use a combination of structure and sequence information to learn what amino acids are likely to form favorable interactions if placed at a given position. In addition to sequence and node embeddings from ProteinMPNN, we also extract directed edge features representing the interaction between the mutated residue pair (Fig. 1B). We formulate our model as a Siamese network by passing the concatenated per-mutation features through a shared prediction head twice, once in each possible order. The raw predicted scores (ΔΔG_AB_ and ΔΔG_BA_) are then symmetrized using a specialized loss function to enforce invariance to the mutation order (details in Methods). We train ThermoMPNN-D on the double mutant subset of the Megascale cDNA proteolysis dataset from Tsuboyama et al.^16^, which we call Megascale-D. Using this scheme, ThermoMPNN-D obtains a high degree of order-invariance, with a Spearman correlation coefficient (SCC) of 0.999 and average bias of 0.003 between ΔΔG_AB_ and ΔΔG_BA_ across the Megascale-D test set.

**Figure 1:**
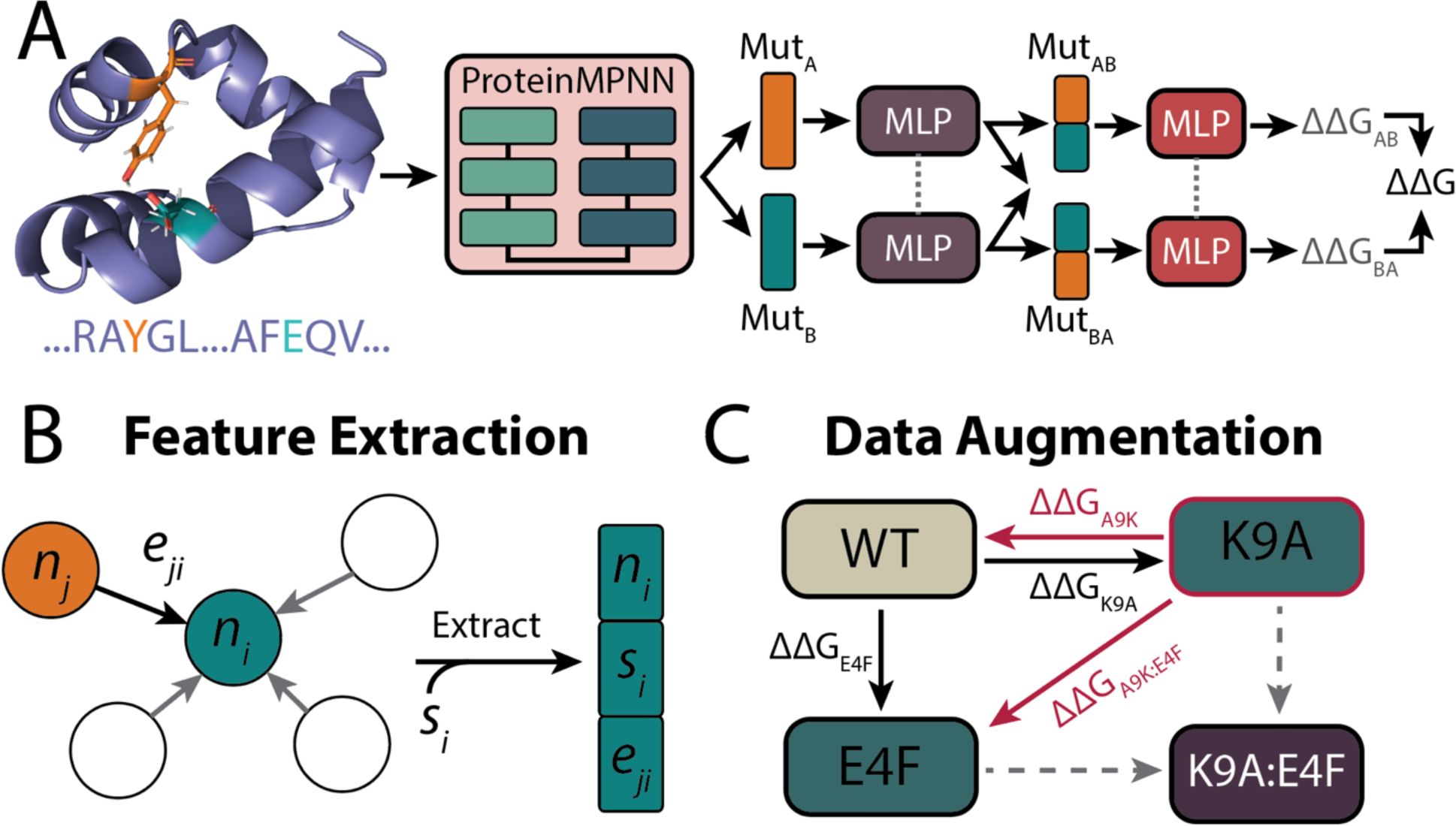
The ThermoMPNN-D modeling framework. A) Schematic of ThermoMPNN-D, a Siamese neural network for predicting double mutant stability changes. Dashed grey lines indicate shared weights. B) Example feature extraction step for hypothetical mutation *i*, in which the node (*n_i_*), sequence (*s_i_*), and edge (*e_ji_*) embeddings are extracted from the protein graph. C) Thermodynamic cycle demonstrating the principle of over-and-back data augmentation. Black arrows denote mutations with a defined ΔΔG in the original dataset, dashed grey arrows indicate mutations missing data, and red arrows indicate mutations defined only via augmentation. The augmented wildtype state is outlined in red.

Training ThermoMPNN-D on the Megascale-D dataset produced reasonable results on the test split of the same dataset (SCC = 0.49 ± 0.01), but it struggled to generalize when tested on an orthogonal test set from the literature, the Protherm double mutant dataset^17^, which we call PTMUL-D (SCC = 0.35 ± 0.03) (Table 1, top section). After examining Megascale-D, we found that, unlike its single mutant counterpart (Megascale-S), it is skewed in several ways. Most notably, mutated residue pairs are typically close in 3D space, often in direct contact via side chain interactions (Fig. 2A, blue bars), with a mean pairwise distance of 3.7Å. Wildtype residue pairs in the dataset also tend to consist of large polar or aromatic groups engaged in strong couplings such as hydrogen bonds and pi-cation interactions (Fig. 2B, blue bars). We hypothesized that training on a dataset with these characteristics may lead to subpar generalizability. To address this issue, we propose a new data augmentation trick which we call over-and-back data augmentation.

**Figure 2:**
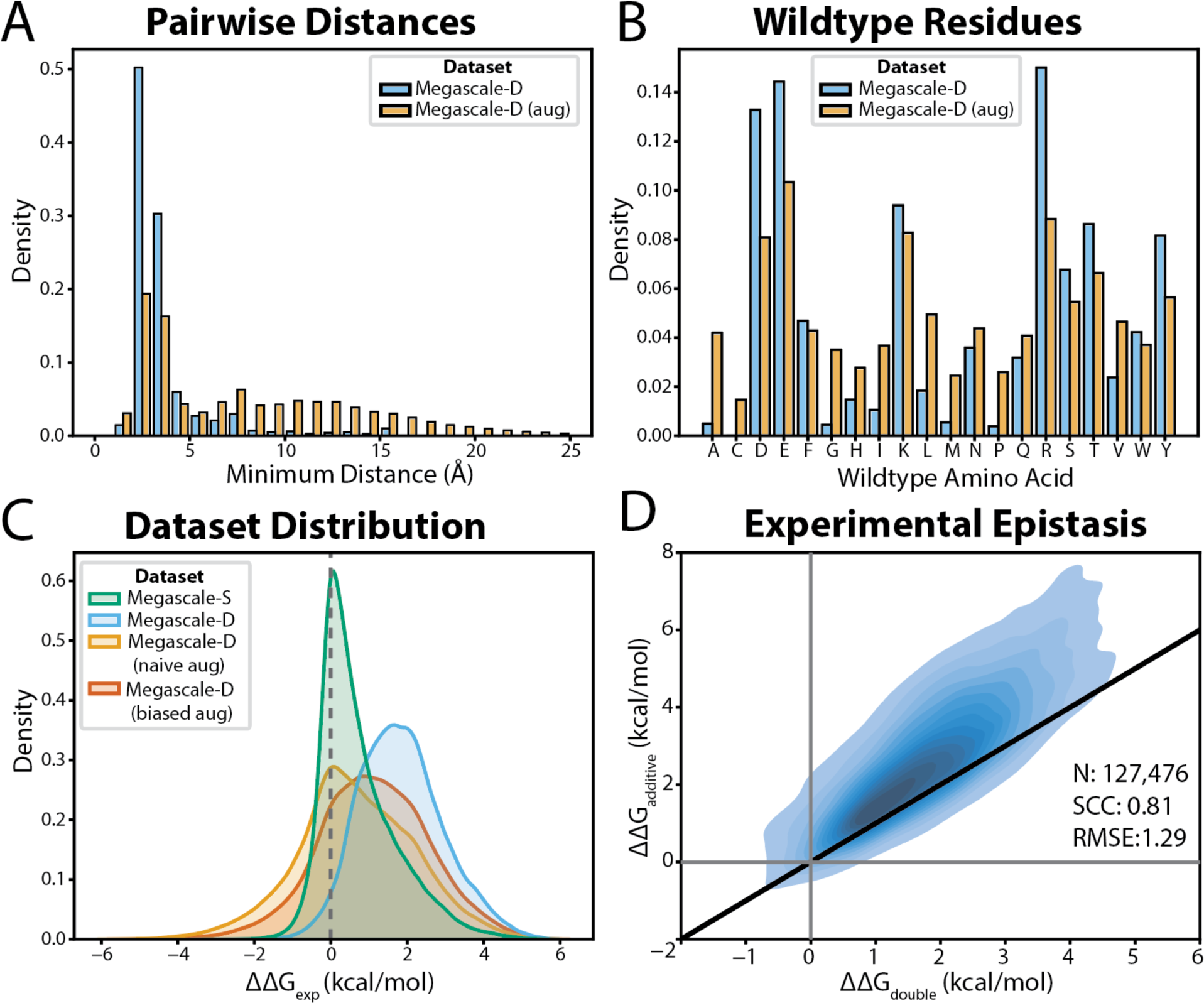
Megascale double mutant (Megascale-D) dataset analysis and augmentation. A) Frequency of mutations stratified by minimum pairwise interatomic distance between mutated residues and B) frequency of wildtype amino acids in the original and augmented Megascale-D. C) Kernel density estimate distributions of Megascale dataset ΔΔG values with and without augmentation. Dashed grey line indicates a theoretical neutral mutation. More positive ΔΔG values indicate more destabilizing mutations. D) Kernel density estimate plot of Megascale-D comparing measured double mutant ΔΔG to the corresponding additive ΔΔG obtained from the sum of the two constituent single mutants. The identity line is shown in black.

**Table 1:** ThermoMPNN-D ablation study results. D.A. stands for data augmentation.

| Trial | Megascale-D |  |  | PTMUL-D |  |  |
| --- | --- | --- | --- | --- | --- | --- |
|  | PCC | SCC | RMSE | PCC | SCC | RMSE |
| **No D.A.** | **$0.54 \pm 0.02$** | $0.49 \pm 0.01$ | **$0.96 \pm 0.01$** | $0.36 \pm 0.04$ | $0.35 \pm 0.03$ | $2.11 \pm 0.02$ |
| **Naïve D.A.** | $0.50 \pm 0.02$ | $0.51 \pm 0.02$ | $1.19 \pm 0.03$ | **$0.55 \pm 0.03$** | **$0.57 \pm 0.02$** | $2.06 \pm 0.04$ |
| **Biased D.A.** | $0.52 \pm 0.02$ | **$0.53 \pm 0.02$** | $1.09 \pm 0.02$ | **$0.55 \pm 0.02$** | **$0.57 \pm 0.02$** | **$1.96 \pm 0.03$** |
| **Siamese** | **$0.52 \pm 0.02$** | **$0.53 \pm 0.02$** | **$1.09 \pm 0.02$** | **$0.55 \pm 0.02$** | **$0.57 \pm 0.02$** | **$1.96 \pm 0.03$** |
| **Max** | $0.50 \pm 0.01$ | $0.52 \pm 0.01$ | $1.15 \pm 0.02$ | $0.50 \pm 0.02$ | $0.54 \pm 0.01$ | $2.05 \pm 0.02$ |
| **Mean** | $0.43 \pm 0.01$ | $0.42 \pm 0.01$ | $1.19 \pm 0.02$ | $0.50 \pm 0.01$ | $0.52 \pm 0.01$ | $2.02 \pm 0.01$ |
| **Sum** | $0.45 \pm 0.01$ | $0.43 \pm 0.01$ | $1.19 \pm 0.03$ | $0.50 \pm 0.01$ | $0.53 \pm 0.01$ | $2.02 \pm 0.02$ |
| **Product** | $0.46 \pm 0.04$ | $0.47 \pm 0.03$ | $1.27 \pm 0.02$ | $0.49 \pm 0.03$ | $0.52 \pm 0.02$ | $2.07 \pm 0.03$ |
| **Baseline** | $0.52 \pm 0.02$ | $0.53 \pm 0.02$ | $1.09 \pm 0.02$ | $0.55 \pm 0.02$ | $0.57 \pm 0.02$ | $1.96 \pm 0.03$ |
| **- Edges** | $0.49 \pm 0.01$ | $0.51 \pm 0.01$ | $1.13 \pm 0.01$ | $0.52 \pm 0.01$ | $0.56 \pm 0.02$ | $2.00 \pm 0.01$ |
| **+ Fine-tune** | $0.47 \pm 0.02$ | $0.48 \pm 0.02$ | $1.15 \pm 0.01$ | $0.55 \pm 0.01$ | $0.59 \pm 0.01$ | $1.96 \pm 0.03$ |
| **+ Ensemble** | **0.54** | **0.55** | **1.07** | **0.57** | **0.59** | **1.95** |

All statistics are reported as mean ± s.d. of triplicate runs, except for the ensemble.

### Over-and-back data augmentation

Our key observation is that every pair of single mutations in a protein are separated from each other by two mutations. To construct an augmented data point (Fig. 1C), we select a single mutant to serve as the wildtype state and invert its experimentally measured ΔΔG_single_ to represent the reverse mutation. We then randomly sample a second single mutant within the same protein, but at a different residue position, and add its experimentally measured ΔΔG_single_ to obtain our final ΔΔG_double_. In this way, we can generate a much larger dataset which more evenly samples the expected distribution in terms of pairwise distance and wildtype amino acid types (Figs. 2A and 2B, orange bars). In doing so, we hoped to enable our model to distinguish between distal, roughly additive mutations and proximal, tightly coupled mutations. After retraining on the augmented dataset, we observed significantly better results on PTMUL-D (SCC = 0.57 ± 0.02) at the cost of a small drop in some Megascale-D metrics (Table 1, top panel). We noticed that this procedure tends to generate a disproportionate fraction of stabilizing double mutants. Since most single mutants are destabilizing, flipping the first ΔΔG_single_ tends to bias the resulting distribution toward lower ΔΔG_double_ values (Fig. 2C, yellow peak). To partially correct for this effect, we implemented a biased sampling procedure to shift the distribution closer to that of the non-augmented Megascale-D dataset (Fig. 2C, orange peak). This adjustment slightly improved both root mean squared error (RMSE) and correlation metrics across both datasets (Table 1, top panel).

### ThermoMPNN-D ablation study

We next tested whether the Siamese aggregation scheme was necessary to achieve strong performance (Table 1, middle panel). We found that this approach obtained better results on both datasets when compared to previously proposed order-invariant aggregation functions such as element-wise summation and averaging. We also experimented with modifying or removing other components of our network (Table 1, bottom panel). We found that removing edge features slightly degraded scores, but not as much as removing the Siamese aggregation. Additionally, we tested fine-tuning ProteinMPNN by unfreezing the weights from the sequence recovery encoder/decoder, which are kept fixed by default. Consistent with the original ThermoMPNN study, fine-tuning the ProteinMPNN weights produced mixed results due to overfitting^10^. A small performance gain was achieved by ensembling three independently trained models, a boost that we do not observe when applied to single mutant ThermoMPNN. We suspect that this is enabled by the randomness introduced by the data augmentation procedure. The final ensembled ThermoMPNN-D predictor achieved SCC values of 0.54 and 0.59 on the Megascale-D and PTMUL-D test sets, respectively.

### Benchmarking ThermoMPNN-D against other double mutant models

We then benchmarked ThermoMPNN-D against existing methods for double mutant ΔΔG prediction from the literature (Fig. 3). To do so, we performed 5-fold cross-validation across the full Megascale-D dataset. We found that ThermoMPNN-D achieved state-of-the-art performance on PTMUL-D, while only recent AlphaFold-based method Mutate Everything obtained comparable performance on Megascale-D when evaluated on matching splits (SCC = 0.55). As a baseline, we also included an additive ThermoMPNN prediction in which we simply added the two predicted ΔΔG_single_ values for comparison to the epistasis-aware prediction of ThermoMPNN-D. To our surprise, this method achieved even better results on Megascale-D (SCC = 0.59), along with similar results on PTMUL-D, depending on the splits used. Intrigued by this finding, we reevaluated each double mutant predictor from the literature by running a similar additive baseline when available (Fig. 3A and 3B, green bars). We found that most methods provide little or no improvement over their additive equivalent when utilized in epistatic mode. The only epistasis-aware methods to provide better scores on both datasets were Rosetta and ESM-1v.

**Figure 3:**
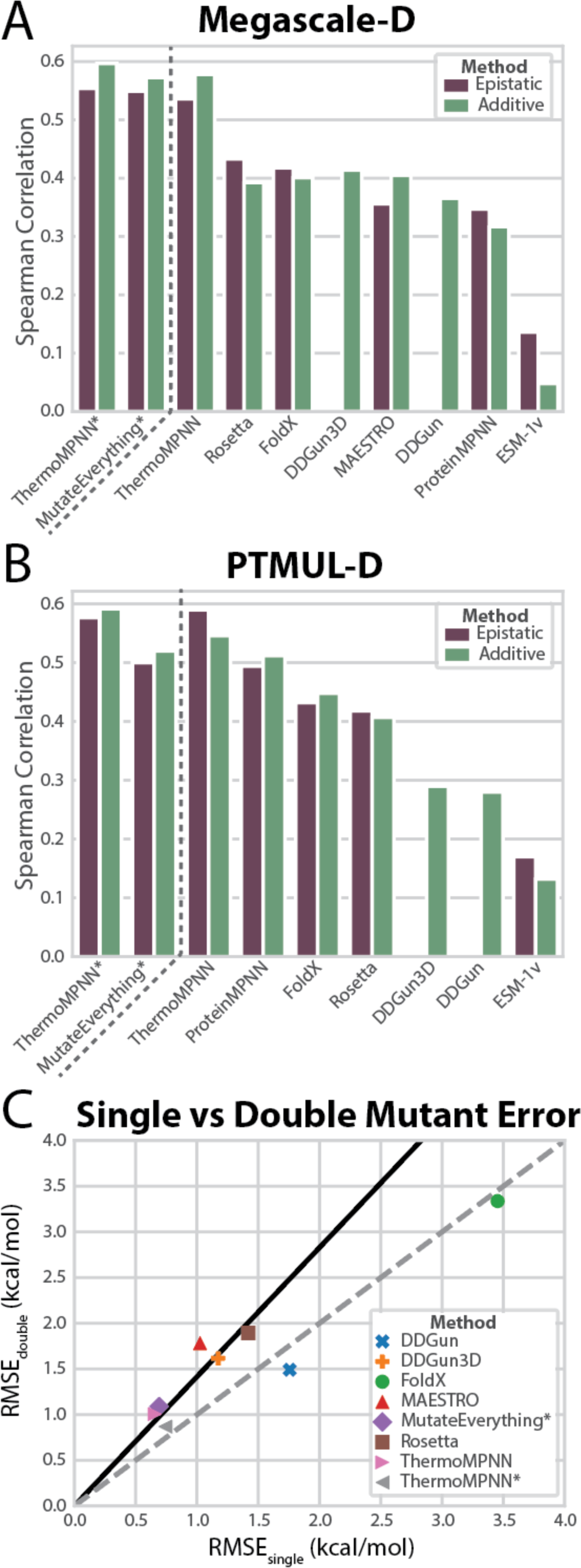
Comparison of ThermoMPNN and selected prior methods for modeling double mutants. A-B) Spearman correlation of selected additive and epistatic methods on the Megascale double mutant dataset (N=127,476) and B) the PTMUL double mutant dataset (N=536). Methods marked with asterisks were retrained and evaluated using different Megascale dataset splits. C) Root mean squared error (RMSE) of selected methods on the Megascale single mutant (x-axis) and double mutant (y-axis) datasets. The identity line is shown in dashed grey, and the theoretical error for a method following naïve additive error propagation behavior is shown in solid black.

To further probe this phenomenon, we evaluated each predictor on Megascale-S for the same set of proteins. We then plotted the single and double mutant error (RMSE) for each method (Fig. 3C). All but two methods had lower error on single mutants, and they closely followed the expected trajectory for the propagation of random additive errors. This indicates that the surveyed methods generally fail to reduce the error on double mutants beyond what would be expected from a purely additive model. The other two methods, FoldX and DDGun, instead followed the identity line, with similar error on single and double mutants.

Since the Megascale dataset includes single and double mutant scans for the same proteins, we can calculate the expected ΔΔG for a particular double mutant assuming an additive model (ΔΔG_additive_). We plotted these values against the measured ΔΔG_double_ for the full Megascale-D dataset (Fig. 2D). Notably, ΔΔG_double_ is highly correlated with ΔΔG_additive_ across the dataset (SCC = 0.81), while the average observed epistatic coupling is −0.9 kcal/mol, indicating that ΔΔG_double_ is typically less destabilizing than would be expected based on the observed ΔΔG_single_. Fitting a linear regression to this dataset produces a y-intercept of 0.62 and a slope of 1.15, indicating that the magnitude of epistatic effects increases with increasing ΔΔG_additive_.

### Deep mutational scan benchmark

We next tested the same predictors on a collection of six deep mutational scans (DMS) gathered from the literature (Table 2). Each DMS dataset consisted of at least 1,000 phenotypic measurements for double mutants gathered in a single study (details in Table 3). Since these assays each measure some proxy of protein fitness rather than stability, we anticipated lower correlations with predicted ΔΔG than on the previous datasets. This was observed in most cases, and the best methods across the full suite of assays were Mutate Everything (additive) and FoldX (epistatic), with average SCC values of 0.40 and 0.39, respectively. Consistent with the prior results, most methods show similar or worse performance in epistatic mode. Only FoldX produced equivalent or better scores across all DMS assays.

**Table 2:** Deep mutational scan benchmark results for selected double mutant prediction methods (additive/epistatic models). The score of the best method on each assay is bolded.

| Model | Spearman Correlation Coefficient |  |  |  |  |  | Mean |
| --- | --- | --- | --- | --- | --- | --- | --- |
|  | avGFP | cgreGFP | ppluGFP2 | amacGFP | His3 | KRas |  |
| **Rosetta**<sup>11</sup> | 0.41/0.42 | 0.37/0.34 | 0.29/0.29 | 0.21/0.21 | 0.26/0.26 | 0.37/0.35 | 0.32/0.31 |
| **FoldX**<sup>31</sup> | 0.46/0.47 | **0.52/0.52** | **0.39/0.39** | **0.38/0.38** | 0.20/0.26 | 0.34/0.34 | 0.38/0.39 |
| **DDGun**<sup>17</sup> | 0.13/-- | 0.32/-- | 0.17/-- | 0.14/-- | 0.14/-- | 0.21/-- | 0.19/-- |
| **DDGun3D**<sup>17</sup> | 0.27/-- | 0.31/-- | 0.18/-- | 0.16/-- | 0.11/-- | 0.24/-- | 0.21/-- |
| **MAESTRO**<sup>32</sup> | 0.26/0.22 | 0.23/0.15 | 0.14/0.08 | 0.11/0.07 | 0.17/0.13 | 0.25/0.26 | 0.19/0.15 |
| **ESM-1v**<sup>33</sup> | 0.00/0.01 | 0.01/0.02 | -0.01/0.02 | -0.01/0.01 | 0.14/0.21 | 0.19/0.20 | 0.05/0.08 |
| **ProteinMPNN**<sup>15</sup> | 0.35/0.36 | 0.23/0.26 | 0.12/0.11 | 0.13/0.12 | 0.18/0.15 | 0.36/0.37 | 0.23/0.22 |
| **ThermoMPNN** | 0.46/0.40 | 0.40/0.24 | 0.21/0.03 | 0.26/0.16 | 0.28/0.24 | 0.37/0.31 | 0.33/0.23 |
| **ThermoMPNN\*** | 0.48/0.44 | 0.40/0.22 | 0.21/0.06 | 0.28/0.18 | **0.29/0.24** | 0.39/0.31 | 0.34/0.24 |
| **Mutate Everything**<sup>19</sup> | **0.53/0.49** | 0.50/0.43 | 0.37/0.30 | 0.32/0.27 | 0.27/0.27 | **0.40/0.36** | **0.40/0.35** |

\* Retrained on cDNA training splits from Ouyang-Zhang et al.<sup>19</sup>

**Table 3:** Summary of curated deep mutational scan assays of double mutants.

| Assay name and source | Abbreviation | Mutations | Phenotype |
| --- | --- | --- | --- |
| GFP_AEQVI<sup>35</sup> | avGFP | 12,777 | Fluorescence |
| D7PM05_CLYGR<sup>36</sup> | cgreGFP | 10,148 | Fluorescence |
| Q6WV13_9MAXI<sup>36</sup> | ppluGFP2 | 15,992 | Fluorescence |
| Q8WTC7_9CNID<sup>36</sup> | amacGFP | 11,260 | Fluorescence |
| HIS7_YEAST<sup>37</sup> | His3 | 1,475 | Enzyme activity |
| RASK_HUMAN<sup>38</sup> | KRas | 22,946 | Expression |

### Stabilizing mutation detection

We next evaluated stabilizing mutation predictions across the Megascale-D and PTMUL-D datasets (Table 4). Stabilizing mutations are particularly difficult to predict, since the vast majority of mutations are typically neutral or destabilizing compared to the wildtype. Indeed, less than 1% of mutations in Megascale-D (n=1,254) fell under the commonly used threshold of ΔΔG ≤ −0.5 kcal/mol. Surprisingly, nearly every predictor showed improvement on both datasets when in epistatic mode. While positive predictive value (PPV) showed mixed results in some cases, all other metrics including Matthews Correlation Coefficient (MCC) consistently favored the epistatic predictors. ThermoMPNN-D achieved the best prediction metrics on the Megascale-D and PTMUL-D datasets, with an MCC of 0.19 and 0.38, respectively, compared to 0.17 and 0.28 for additive ThermoMPNN. When evaluated on the cDNA2 test split of Megascale-D, Mutate Everything (epistatic) outperforms ThermoMPNN-D (MCC = 0.27 vs 0.15), but the latter is more effective on the PTMUL-D dataset when trained on the same splits (MCC = 0.38 for ThermoMPNN-D vs 0.33 for Mutate Everything). We observe a significant discrepancy in stabilizing mutation scores between PTMUL-D and Megascale-D, with nearly all methods producing significantly better metrics on PTMUL-D in both additive and epistatic mode.

**Table 4:** Stabilizing mutation detection metrics for selected prediction methods (additive/epistatic models). The score of the best method on each metric is bolded.

|  | **Megascale-D (n=1,254)** |  |  |  | **PTMUL-D (n=111)** |  |
| --- | --- | --- | --- | --- | --- | --- |
| **Model** | **MCC** | **PPV** | **DetPr<sub>30</sub>** | **nDCG<sub>30</sub>** | **MCC** | **PPV** |
| **Rosetta**<sup>11</sup> | 0.11/0.15 | 0.05/0.09 | 0.07/0.12 | 0.15/0.20 | 0.29/0.29 | 0.48/0.42 |
| **FoldX**<sup>31</sup> | 0.13/0.14 | 0.04/0.04 | 0.07/0.08 | 0.16/0.16 | 0.22/0.24 | 0.38/0.36 |
| **DDGun**<sup>17</sup> | 0.12/-- | 0.04/-- | 0.10/-- | 0.18/-- | 0.22/-- | 0.50/-- |
| **DDGun3D**<sup>17</sup> | 0.13/-- | 0.05/-- | 0.08/-- | 0.17/-- | 0.17/-- | 0.47/-- |
| **MAESTRO**<sup>32</sup> | 0.15/0.14 | 0.04/0.03 | 0.09/0.09 | 0.13/0.17 | --/-- | --/-- |
| **ESM-1v**<sup>33</sup> | 0.02/0.03 | 0.01/0.02 | 0.03/0.05 | 0.05/0.12 | 0.07/0.09 | 0.30/0.31 |
| **ProteinMPNN**<sup>15</sup> | 0.07/0.10 | 0.06/0.05 | 0.07/0.09 | 0.17/0.19 | 0.30/0.33 | 0.51/0.49 |
| **ThermoMPNN** | 0.17/**0.19** | **0.13/0.13** | 0.20/**0.22** | 0.31/**0.35** | 0.29/**0.37** | 0.49/**0.57** |
|  | **cDNA2 test (n=198)** |  |  |  | **PTMUL-D (n=111)** |  |
| **ThermoMPNN\*** | 0.10/0.15 | **0.29/0.20** | 0.10/0.17 | 0.11/0.22 | 0.34/**0.38** | **0.58/0.54** |
| **Mutate Everything**<sup>19</sup> | 0.26/**0.27** | 0.12/0.11 | 0.24/**0.30** | 0.35/**0.43** | 0.33/0.33 | 0.46/0.44 |

\* Re-trained on cDNA training splits from Ouyang-Zhang et al.<sup>19</sup>

## Discussion

This study was motivated by the hypothesis that a network designed to explicitly model double point mutations could provide better ΔΔG predictions than a naïve model assuming additive mutational effects. To test this hypothesis, we developed ThermoMPNN-D, which uses a Siamese aggregation scheme and extensive data augmentation to leverage extensive mutagenesis data and enforce helpful inductive biases such as the distance dependence of epistatic interactions and mutation order invariance. Through rigorous benchmarking, we found our initial hypothesis was not always correct, as ThermoMPNN-D and other double mutant predictors nearly all achieved similar or worse overall results than their additive counterparts when evaluated by full-dataset correlation coefficients. However, epistasis-aware predictors including ThermoMPNN-D enabled improved prediction of stabilizing double mutations, which are critically important for protein design applications.

Our study is one of the first to utilize the double mutant subset of the Megascale cDNA proteolysis dataset recently published by Tsuboyama et al.^18^, which we call Megascale-D. As such, it is important to note that models trained solely on Megascale-D proved unable to generalize to unseen datasets. To address this issue, we introduce a novel data augmentation technique, over-and-back augmentation, which may be considered as an extension of the recently introduced thermodynamic permutation technique^9^ for sampling double mutations. The other extant study utilizing the Megascale-D dataset also chose to expand their training dataset by pre-training on Megascale-S^19^, although they did not evaluate a model trained only on Megascale-D. Taken together, these findings raise the question: what constitutes a representative double mutant landscape for modeling purposes? While exhaustive single mutant scans are now feasible for small proteins, enumeration of double mutations remains challenging due to the exponential increase in scale. With this in mind, we contend that data augmentation is an attractive strategy to expand the pool of double mutant data to better capture the full mutational landscape. To enable further development of data augmentation protocols, we make readily available our full dataset of 340,000 modeled mutant structures and Rosetta energies.

Most other protein stability models are limited to predicting single point mutations, while even those with multiple mutation functionality have rarely been benchmarked against an appropriate additive baseline. Still, a few previous studies provide evidence to corroborate our findings. Consistent with our observations, Ouyang-Zhang et al. find that the epistatic version of Mutate Everything behaves similarly to ThermoMPNN-D, in that its overall regression metrics are similar or worse compared to its additive equivalent despite showing improved prediction of stabilizing double mutations^19^. We also found that epistasis-aware models were often better performing on certain datasets but worse on others. This is consistent with prior works which find that epistatic terms derived from coevolutionary models are only beneficial for around 2/3 of tested proteins^20,21^, with factors such as MSA depth and assay design suggested as possible explanations.

We anticipated that predicting ΔΔG for double mutations would be more difficult than for single mutations. This was generally observed, as top predictors including ThermoMPNN obtained an SCC below 0.60 on Megascale-D, while the top reported score^10^ on Megascale-S is around 0.75. As expected, we also observe a lower success rate on stabilizing mutations, as ThermoMPNN obtains a state-of-the-art PPV of 0.13 and 0.29 on different splits of Megascale-D compared to 0.45 on Megascale-S^10^. Double mutant data is less abundant than single mutant data, which makes benchmarking more prone to random variance. To alleviate this issue, we employ DMS data to supplement our stability datasets and cross-validate across all available data, which enabled evaluation of >125,000 stability measurements and >74,000 DMS measurements gathered on double mutations. Future work includes benchmarking and model development on higher-order (3+) mutation datasets, which face even greater limitations in data availability and evaluation.

Epistasis is a complex phenomenon in which both global (per-protein) and local (per-mutation) effects can influence variant fitness^22^, and their relative importance can vary by fitness level and biological context^23^. With this in mind, several avenues for future work may offer potential for improvement. The pre-training schemes underpinning many recent models may be redesigned to explicitly learn patterns of epistatic interaction rather than autoregressive or one-shot decoding schemes. Model architecture may also be improved either by separating energetic contributions from individual and pairwise residue terms, such as with a Potts model^24^, or by incorporating latent variables to represent global nonlinearities^25^. Recent efforts to model protein fitness with epistasis-aware neural networks^26,27^ may serve as a starting point for future protein stability models. However, these methods tend to require parameterization with initial DMS data for the target protein, so it remains to be seen how well they can generalize to novel proteins.

## Methods

### ThermoMPNN-D architecture

ThermoMPNN-D (Fig. 1A) was implemented as an extension of the ThermoMPNN framework^10^, which uses sequence recovery model ProteinMPNN as a feature extractor^15^. All experiments used the ProteinMPNN model trained with 0.2Å backbone noise, and ProteinMPNN weights were kept frozen during training unless otherwise stated. To represent each mutation, we extracted the node representation *n_i_* for the mutated position from the molecular graph held in the last two decoder layers of ProteinMPNN. We also retrieved the directed edge representation *e_ji_* connecting from the other mutated residue to the residue of interest (Fig. 1B). If no such edge existed (i.e., the mutations are not within 48 nearest neighbors), a zero vector was substituted as the edge representation. We subtracted the sequence embedding of the wildtype and mutant amino acids to obtain a sequence representation *s_i_*. The node, edge, and sequence representations were concatenated, and each mutation vector was then passed through a shared MLP to aggregate and downsample to 128 dimensions. The mutation features were then concatenated in both AB and BA order, and each permutation was passed through another shared MLP to produce raw predictions ΔΔG_AB_ and ΔΔG_BA_, which were averaged to obtain a final ΔΔG.

### ThermoMPNN-D training procedure

ThermoMPNN-D includes 116,000 trainable parameters, which were trained for up to 100 epochs using the Adam optimizer with an initial learning rate of 10^−5^ and a batch size of 256 mutations. Dropout (p=0.1) and LayerNorm were used on all fully connected layers. Learning rate decay and early stopping was conditioned on validation set mean squared error (MSE). Training used a custom loss function inspired by antisymmetric single mutant predictor ACDC-NN^28^ and applied to the raw predictions ΔΔG_AB_ and ΔΔG_BA_:

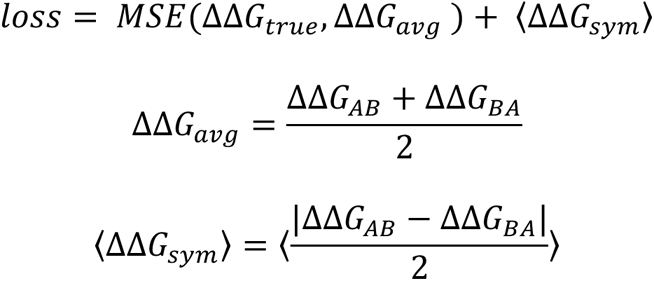

A non-Siamese model was built to test other aggregators (Table 1, middle panel). This model used the same featurization scheme, but after downsampling, mutation embeddings were aggregated instead of concatenated and passed once through the final MLP. Fine-tuning ProteinMPNN was implemented by unfreezing all layers with a separate learning rate, which was selected via parameter sweep (10^-6^ gave the best results). Ensembling was implemented by averaging the predicted ΔΔG from three independently trained models with different random seeds for training and data augmentation.

### Over-and-back data augmentation

For each single mutant in the Megascale training set, the modeled mutant structure was obtained using Rosetta^11^. The second mutation was sampled stochastically from all possible single mutations that a) shared the same PDB ID and b) did not share the same amino acid position. To bias sampling toward more destabilizing ΔΔG values, the ΔΔG_single_ values for the whole dataset were used to obtain a weighted sampling probability (P) as follows:

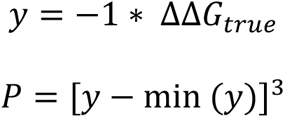

This distribution was normalized for each individual mutation. Augmented datasets were sampled once at the beginning of training and randomly shuffled after every epoch.

### Dataset splits and curation

For the ThermoMPNN-D ablation study, we obtained the Megascale dataset reported in Tsuboyama et al.^18^ from its Zenodo repository^16^, following the splitting procedure previously described for ThermoMPNN^10^, with the following modifications. We removed any homologues (>25% sequence similarity) to proteins in the PTMUL dataset. Second, we trained on double mutants with defined ddG_ML values. After removing duplicate data points, we obtained a training/validation/test split of 85,253/10,282/18,574 mutations across 90/17/20 proteins.

For the double mutant model benchmarks, we used the full Megascale dataset and evaluated ThermoMPNN using 5-fold cross-validation split by sequence similarity, as previously described. To compare additive and epistatic models, we matched single and double mutant measurements and dropped any double mutants without valid single mutant data, resulting in 127,476 double mutations across 153 proteins. The Protherm multiple mutation (PTMUL) dataset introduced in the DDGun paper^17^ and re-curated for Mutate Everything^19^ was used after dropping higher-order (3+) mutation measurements, resulting in 536 mutations across 83 proteins (PTMUL-D). Since Mutate Everything was trained on different splits of the Megascale dataset, we retrained and reevaluated ThermoMPNN using their training/test splits, which they denote “cDNA2”, resulting in a test set of 22,913 mutations across 18 proteins. For the single vs double mutant error calculation, we used the full single mutant Megascale dataset (Megascale-S), which contained 271,231 mutations across 298 proteins.

We curated deep mutational scanning (DMS) datasets from the ProteinGym benchmark^29^. We selected DMS datasets with >1000 double mutations and endpoints that might serve as reasonable proxies for thermodynamic stability. From this pool, we eliminated assays overlapping with the Megascale dataset and those without a high-confidence AlphaFold model or crystal structure. We were then left with six assays, which are summarized in Table 3.

### Literature model benchmarking

For the Rosetta benchmark, we adapted a previously published *RosettaScripts* point mutation protocol^11^ for use on double mutations by applying constraints to all residues nearby to either residue. To convert REU into approximate kcal/mol units, we divided all energy values by 2.9, as recommended for the ref2015 score function^30^. FoldX was downloaded under an academic license (https://foldxsuite.crg.eu), and predictions were obtained by running RepairPDB on all input structures, followed by PositionScan for single mutants or additive predictions and BuildModel for epistatic predictions^31^. MAESTRO^32^ was downloaded from its website (https://pbwww.services.came.sbg.ac.at), while DDGun/DDGun3D^17^ (https://github.com/biofold/ddgun), ESM-1v^33^ (https://github.com/facebookresearch/esm), ProteinMPNN^15^ (https://github.com/dauparas/ProteinMPNN), and Mutate Everything^19^ (https://github.com/jozhang97/MutateEverything) were obtained from their respective GitHub repositories.

ProteinMPNN zero-shot predictions were obtained by masking out the mutated residue(s) and calculating the difference in negative log-likelihood between the mutant and wildtype residues. For the ESM zero-shot predictions, we used an ensemble of five ESM-1v (650M, UR90S) models with the masked-marginals scoring method, as recommended^33^. To obtain epistatic predictions for ProteinMPNN and ESM-1v, both mutated residues were masked prior to inference, while the additive predictions masked each residue individually.

### Theoretical error calculation

We calculated the theoretical error for double mutant predictions as follows:

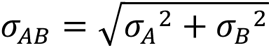

Where *σ_A_* and *σ****_B_*** are the single mutant prediction errors (in RMSE) for mutation A and B, and *σ_AB_*, is the theoretical error for double mutants. Note that this model assumes that single mutant errors are randomly distributed and uncorrelated.

### Stabilizing mutation metrics

To evaluate stabilizing mutation predictions (Table 4), we primarily use the Matthews correlation coefficient (MCC), which is widely accepted as a robust holistic measure of classifier accuracy on unbalanced datasets^34^. Following the convention from Ouyang-Zhang et al.^19^, we calculate MCC across the full dataset using a threshold of 0 kcal/mol. For the remaining metrics, we use the definition that mutations with ΔΔG ≤ −0.5 kcal/mol are stabilizing. This resulted in 1254, 111, and 198 stabilizing mutations for the Megascale-D, PTMUL-D, and cDNA2 test datasets, respectively.

We calculate the positive predictive value (PPV) across each full dataset, while detection precision (DetPr) and normalized discounted cumulative gain (nDCG) are calculated separately for each protein and averaged. To calculate these last two metrics, the mutations for a given protein are sorted by predicted ΔΔG, and the top K mutations are selected (K=30 in this study). The DetPr represents the fraction of top-30 mutations that are measured to be truly stabilizing, while nDCG is a more complicated measure of how highly the model ranks the best 30 mutations.

## Code Availability

ThermoMPNN-D trained model weights and code are available at https://github.com/Kuhlman-Lab/ThermoMPNN-D.

## Data Availability

The full Megascale dataset can be obtained from its Zenodo repository^16^, while the full ProteinGym datasets are available at https://proteingym.org^29^ and the full PTMUL dataset is available at https://github.com/jozhang97/MutateEverything. The curated Megascale, PTMUL-D, and DMS double mutant datasets and splits used in this study are available on Zenodo at https://doi.org/10.5281/zenodo.13345274. Modeled single mutant structures and energies obtained using Rosetta for the full Megascale dataset are available in the same repository.

## Supplementary material description

N/A

## Conflict of interest statement

The authors have no relevant conflicts of interest to declare.

## Acknowledgements

This work was supported by NIH grant R35GM131923 (B.K.) and NSF fellowship DGE-2040435 (H.D.). H.D. acknowledges support by a Pre-doctoral Fellowship from the American Foundation for Pharmaceutical Education. This work utilized the resources of the UNC Longleaf high-performance computing cluster. The authors would like to thank Dr. Pranam Chatterjee for his advice regarding protein language models and Jeffrey O. Zhang for his assistance with the Mutate Everything platform.

## Notes

https://github.com/Kuhlman-Lab/ThermoMPNN-D

https://zenodo.org/records/13345274

### Competing Interest Statement

The authors have declared no competing interest.

